# Snurportin-1 maintains muscle niche integrity and myogenic progenitor homeostasis

**DOI:** 10.64898/2026.09.24.749599

**Authors:** Hilal Pırıl Saraçoğlu, Marwan Nashabat, Duygu Naz Kutlu-Bilgili, Burak Sarıbaş, Elanur Yılmaz, Şeymanur Tur, Hülya Kayserili, Emre Yakşi, Nasrinsadat Nabavizadeh, Nathalie Escande-Beillard

## Abstract

Loss-of-function variants in *SNUPN*, encoding the nuclear import factor Snurportin-1 (SPN1) required for spliceosomal small nuclear ribonucleoprotein (snRNP) transport, cause a recently described form of limb-girdle muscular dystrophy (LGMD). However, the role of SPN1 in skeletal muscle homeostasis remains poorly understood, in part due to the lack of a suitable *in vivo* model. Here, we generated a zebrafish *snupn* loss-of-function model that recapitulates key features of the skeletal muscle phenotype observed in patients. Mutant larvae developed severe locomotor impairment by 6 days post-fertilization (dpf), accompanied by sarcomeric disorganization and impaired muscle fiber integrity.

Transcriptomic profiling at 6 dpf revealed widespread alternative splicing and transcriptional dysregulation, with prominent alterations in extracellular matrix and basement membrane components, together with upregulation of stress- and inflammation-associated genes. Notably, these late-stage abnormalities were preceded by disruption of the muscle progenitor population at 2 dpf, with reduced Pax7^+^ progenitor abundance and myogenic gene expression together with altered muscle differentiation and organization.

Together, these findings identify SPN1 as a key regulator of skeletal muscle homeostasis linking RNA processing to extracellular niche integrity and myogenic progenitor maintenance. This zebrafish model provides an *in vivo* platform for dissecting LGMD-associated disease mechanisms and developing therapeutic strategies aimed at restoring muscle function and regenerative capacity.

## Introduction

Biallelic loss-of-function variants in *SNUPN* have recently been identified as the cause of a novel form of limb-girdle muscular dystrophy (LGMD; MIM#620793), expanding the genetic landscape of these heterogeneous disorders ^1–3^. *SNUPN* encodes Snurportin-1 (SPN1), a nuclear import receptor required for the transport and maturation of spliceosomal small nuclear ribonucleoproteins (snRNPs) into the nucleus. SPN1 recognizes the trimethylguanosine (TMG) cap of spliceosomal snRNAs and, with importin-β, mediates their nuclear import, enabling their maturation and assembly into functional snRNPs required for pre-mRNA splicing. This pathway is functionally connected to the survival motor neuron (SMN) machinery, which promotes the assembly and maturation of spliceosomal snRNPs ^4–8^.

SPN1-dependent snRNP trafficking is therefore essential for maintaining snRNP homeostasis and proper RNA processing, yet how disruption of this ubiquitous cellular pathway leads to tissue-specific pathology predominantly affecting skeletal muscle remains unclear ^4–8^. Recent work has shown that *SNUPN* variants associated with spinocerebellar atrophy (SCA) are allelic to those causing LGMD, yet result in distinct phenotypes ^9^. Notably, a knock-in mouse model recapitulated the cerebellar phenotype but exhibited minimal muscle involvement, suggesting that current mammalian models may not adequately capture the primary muscular defects observed in LGMD patients.

Our previous studies using patient samples, and gene-edited cellular models revealed that SPN1 deficiency profoundly disrupts RNA processing and muscle integrity, with splicing defects identified in patient muscle and fibroblasts, together with extracellular matrix (ECM) remodeling, sarcolemma instability, defects in the dystrophin–glycoprotein complex (DGC), and cytoskeletal disorganization ^2,3^. However, how these molecular and cellular defects converge to disrupt muscle homeostasis *in vivo* remains poorly understood.

Skeletal muscle homeostasis critically depends on the coordinated interaction between myofibers, the ECM, and muscle stem and progenitor cells that contribute to muscle growth and regeneration^10–12^. The ECM and basement membrane not only provide structural support but also establish the niche required for the maintenance, activation, and differentiation of muscle stem and progenitor cells ^13,14^. While disruption of ECM–cytoskeleton interactions is a hallmark of many muscular dystrophies (MDs) ^15,16^, the impact of defective SPN1 and RNA processing on the integrity of the muscle niche and progenitor cell homeostasis remains unknown.

Here, we developed a CRISPR/Cas9-mediated zebrafish model of *snupn* deficiency. Zebrafish are an established vertebrate model for studying muscular dystrophies, with conserved muscle organization and a well-defined program of post-embryonic myogenesis driven by Pax7⁺ myogenic progenitor and satellite-like cells ^17–20^. Notably, zebrafish possess a single *snupn* ortholog, providing a genetically tractable system to directly investigate the consequences of *SNUPN* deficiency *in vivo*.

Using an integrated approach combining morphological, functional, transcriptomic, and cellular analyses, we demonstrate that *snupn* deficiency induces a multi-level impairment of muscle homeostasis recapitulating key features of human *SNUPN*-associated LGMD. Mechanistically, we identify a pathogenic axis linking defective RNA processing to disruption of the muscle niche, impaired muscle progenitor homeostasis, and ultimately progressive muscle degeneration. Furthermore, this mutant model provides a platform to further dissect MD associated disease mechanisms and explore therapeutic strategies targeting muscle degeneration and impaired regeneration^21^.

## Result

### Modeling *snupn* loss-of-function in zebrafish

To define the temporal expression pattern of *snupn* during zebrafish development, we quantified transcript levels from 3.5 to 72 hours post-fertilization (hpf). Quantitative PCR (qPCR) analysis revealed high *snupn* transcript levels during early blastula stages, consistent with strong maternal deposition, followed by a rapid decline during gastrulation and stabilization at lower levels by 24 hpf. This profile closely paralleled that of the maternally expressed gene *nanog* (Fig. 1a). Public transcriptomic datasets further confirmed strong early expression transitioning to a moderate and ubiquitous expression across germ layers, including mesodermal and neuronal tissues with sustained expression in muscle cell populations at later stages (Fig. S1a-b) ^22,23^. The evolutionary conservation of *SNUPN*, the absence of a duplicated *snupn* gene in zebrafish, and its relevance to muscle biology supported the use of zebrafish to model *SNUPN* deficiency (Fig. S1c). We therefore generated a CRISPR/Cas9-mediated mutant line targeting exon 3 of *snupn* (Fig. 1b). Screening identified an 8-bp insertion (c.220_221insACTGGAGC) causing a frameshift and premature stop codon (p.Ser75Trpfs*41), predicted to produce a truncated and likely unstable protein, consistent with a strong loss-of-function allele (Fig. 1b).

**Figure 1:**
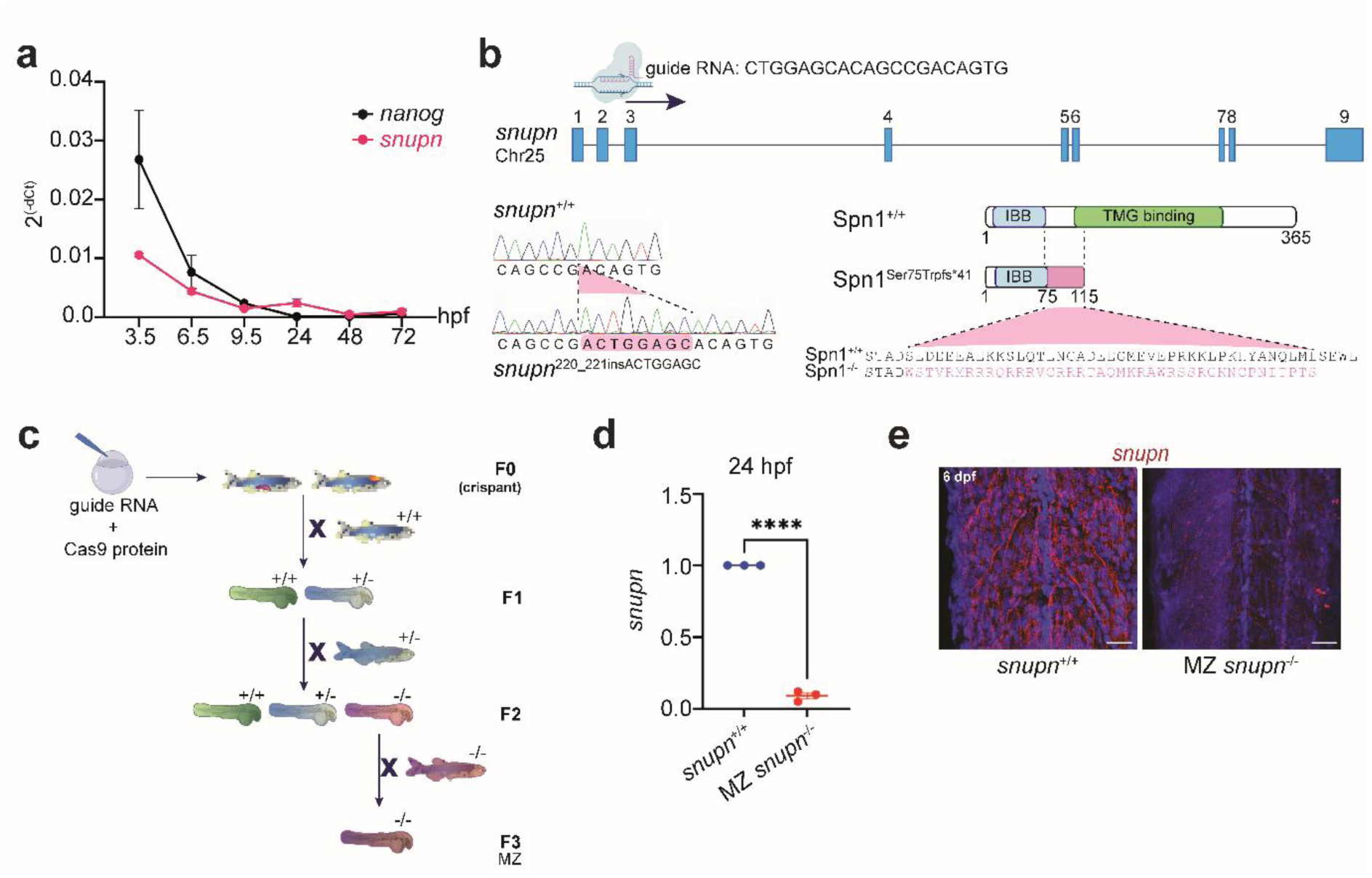
Generation and validation of the MZ *snupn^−/−^* zebrafish line. **a**, Temporal expression profile of *snupn* and *nanog* mRNA during zebrafish embryonic development, as determined by qPCR. hpf: hour post fertilization. Data are normalized to 18S. **b,** Schematic representation of the CRISPR/Cas9 strategy targeting exon 3 of *snupn* to generate the *snupn* mutant line. The guide RNA (gRNA) target site and PAM sequence are indicated. Sequence chromatograms of wild-type (WT) and maternal-zygotic (MZ) *snupn*^−/−^ embryos showing an 8-bp insertion (c.220_221insACTGGAGC) leading to a frameshift and premature stop codon (p.Ser75Trpfs*41). **c**, Guide RNA was co-injected with Cas9 protein into one-cell-stage embryos. Genotyping of F0, F1, F2 and F3 generations identified mosaic crispant, heterozygous (+/−), homozygous (−/−) and MZ mutant lines respectively. **d,** Relative *snupn* transcript levels measured by qPCR in 24 hpf MZ *snupn*^−/−^ mutants compared to WT siblings. Data represent mean ± SEM; n=3 independent experiments,****p<0.0001. **e,** Fluorescent whole-mount Hybridization Chain Reaction (HCR) detecting *snupn* transcripts in WT and MZ *snupn*^−/−^ larvae at 6 days post-fertilization (dpf). Scale bar: 25 µm.

To eliminate maternal *snupn* contribution, homozygous *snupn*^⁻/⁻^ fish were incrossed to generate maternal-zygotic (MZ) mutants (Fig. 1c). qPCR confirmed a significant reduction of *snupn* mRNA levels in both larvae and adult muscle tissues of MZ *snupn*^−/−^ mutants compared to WT siblings (Fig. 1d and S1d). Fluorescent whole-mount Hybridization Chain Reaction (HCR) analysis further validated the robust expression of *snupn* in WT truncal muscle region at 6 dpf and its marked reduction in MZ *snupn*^−/−^ mutant line (Fig. 1e). Together, these results establish a robust genetic model of *snupn* deficiency for functional characterization.

### *snupn* deficiency disrupts skeletal muscle architecture in zebrafish

We next investigated the muscle phenotype of MZ *snupn*⁻/⁻ zebrafish by conducting a series of morphological, histological, and ultrastructural analyses. Although s*nupn* deficiency did not significantly affect overall body length at 7 dpf (Fig. S2a), MZ *snupn*^−/−^ zebrafish exhibited clear abnormalities in muscle architecture and integrity compared to WT siblings.

Birefringence assays revealed a marked reduction in muscle fiber brightness in mutants, indicating impaired fiber integrity and alignment (Fig. 2a). Morphometric measurements at 5 dpf revealed a significant decrease in myotome thickness and increased myotome junction V-shaped angles in MZ *snupn*^−/−^ larvae, further supporting disrupted muscle architecture (Fig. 2b).

**Figure 2:**
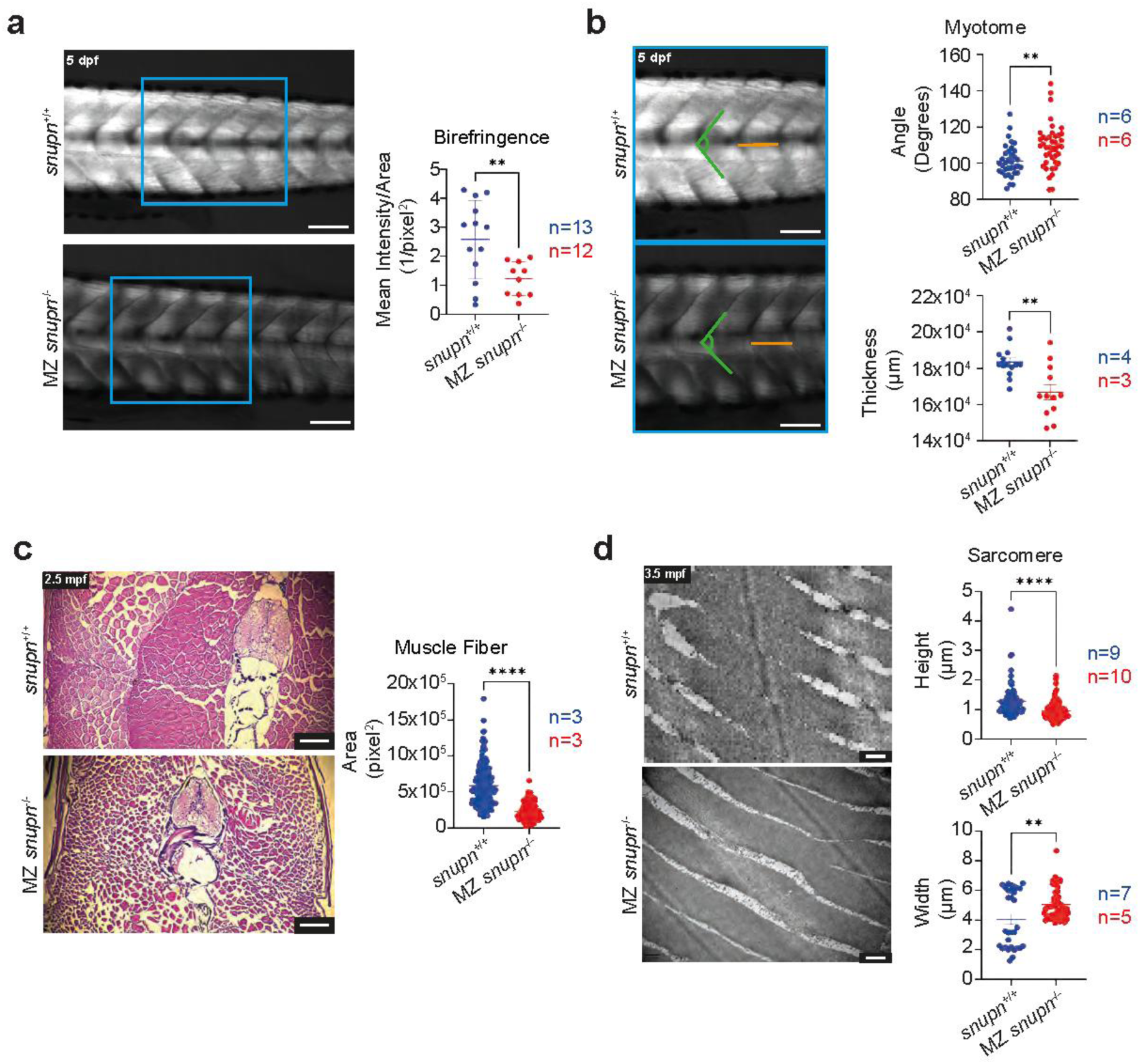
MZ *snupn*^−/−^ larvae display dystrophic muscle phenotype. **a,** Birefringence images showing clear reduction in muscle fiber brightness in 5 dpf MZ *snupn*^−/−^ larvae compared to WT and quantification confirmed significant reduction of mean intensity of birefringence signal, Scale bar: 100 µm. Data represent mean ± SEM, n= 10-13 larvae/genotype, **p=0.0083. **b,** Quantitative analysis at 5 dpf showing reduced myotome thickness and increased myotome junction angle in MZ *snupn*^−/−^ larvae compared to WT. Representative images are shown with orange and green lines indicating thickness and angle measurements, respectively. Scale bar: 100 µm. Data represent mean ± SEM, n= 4-6 larvae/genotype, **p=0.0013 (top) and **p=0.0015 (bottom). **c**, Histological staining of juvenile muscle tissue with Hematoxylin and Eosin (H&E) showing smaller fiber sizes in MZ *snupn*^−/−^ compared to WT controls. Graph showing quantification of muscle fiber area, confirming a significant reduction in the mutant line. Scale bar: 100 µm. Data represent mean ± SEM, n= 3 larvae/genotype, ****p<0.0001. **d,** Electron microscopy of adult muscle tissue reveals severe ultrastructural defects in MZ *snupn*^−/−^ compared to WT controls, including thin muscle fibers and degenerated sarcoplasmic reticulum (SR). Graph showing quantification of height and width of sarcomeres, Scale bars: 500 nm. Data represent mean ± SEM; n= 82-153 sarcomeres from 9-10 images, ****p<0.0001 (top; height) and n= 31-48 sarcomeres from 5-7 images, **p=0.0038 (bottom; width).

Consistently, hematoxylin and eosin (H&E) staining of muscle cross-sections demonstrated a significant overall reduction in muscle fiber area in juvenile mutants compared to WT controls, a hallmark of muscular dystrophy (Fig. 2c). At the ultrastructural level, transmission electron microscopy (TEM) uncovered severe abnormalities, including muscle fiber thinning, degeneration of the sarcoplasmic reticulum (SR), and disrupted mitochondrial morphology in adult MZ mutant line (Fig. 2d; Fig. S2b).

At the molecular level, qPCR analysis demonstrated reduced expression of key components of the dystrophin–glycoprotein complex (DGC), including *lama1* and *dag1*, in mutant larvae and adult muscle, respectively, recapitulating defects observed in *SNUPN-*associated LGMD ^2^ (Fig. S2c).

Together, these findings demonstrate that deficient *snupn* in zebrafish recapitulates key structural and molecular features of *SNUPN*-associated muscular dystrophy.

### *snupn* deficiency impairs motor locomotion in zebrafish

To determine the functional consequences of *snupn* deficiency on motor output, we quantified locomotor behavior in zebrafish larvae using a custom designed behavioral assay, where we can track animals’ swim position, swim speed at rest and response to visual stimuli ^24,25^. MZ *snupn*^−/−^ larvae exhibited markedly disorganized and irregular swimming trajectories, indicative of impaired locomotion (Fig. 3a). Next, we investigated animals’ locomotor responses to light-to-dark transitions ^26^. Mutants’ locomotor responses to light-dark transitions were significantly reduced under both light-on and light-off conditions (Fig. 3b; Fig. S3a-d). All together these results revealed that MZ *snupn*^−/−^ larvae exhibited impairment in spontaneous and stimulus-evoked locomotion.

**Figure 3:**
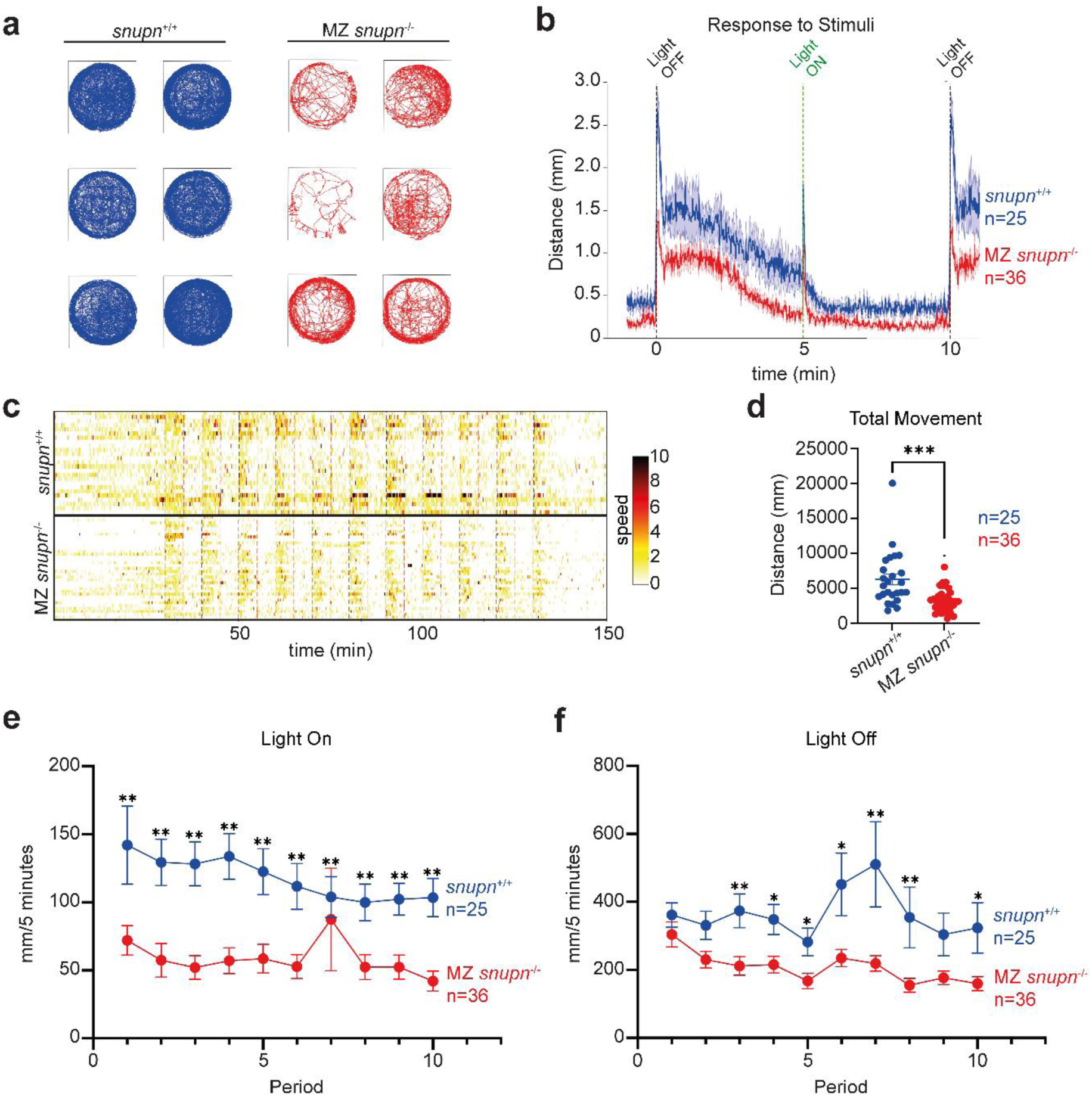
Locomotor activity is reduced in MZ *snupn*^−/−^ larvae. **a,** Representative individual swimming trajectories of 6 dpf larvae, shown in blue for *snupn*^+/+^ and red for MZ *snupn*^−/−^, recorded over a 2.5-hour experimental period, including acclimatization, alternating light-on and light-off phases, and the final recovery period. n=25-36 larvae/genotype. **b,** Graph showing reduction in total distance traveled by larvae at 6 dpf covering period of light on and off of 5 min each. n=25-36 larvae/genotype. **c,** Heatmaps illustrating locomotor activity in WT and MZ *snupn*^−/−^ larvae. The color scale represents movement speed, with darker colors indicating increased activity. **d,** Quantification of total distance traveled per individual larvae, demonstrating significantly reduced locomotor activity in MZ *snupn*^−/−^ larvae. Data represent mean ± SEM, n=25-36 larvae/genotype, ***p=0.0001. **e,f,** Distance traveled within 5-minute intervals during **e**, light-on and **f,** light-off conditions. Data represent mean ± SEM, n=25-36 larvae/genotype, *p<0.05, **p<0.005.

At 6 dpf, mutant larvae demonstrated a clear hypokinetic phenotype, characterized by prolonged immobility, short and discontinuous swim bouts consistent with muscle weakness (Fig. 3c). This was accompanied by a marked reduction in total distance traveled (Fig. 3d). Consistently, average swimming velocity was significantly reduced in MZ *snupn*^−/−^ larvae compared to WT controls, independent of illumination conditions (Fig. 3e-f).

Collectively, these results establish that *snupn* deficiency translates into significant *in vivo* functional impairment, substantiating its critical role in maintaining muscular performance and reinforcing its relevance to human *SNUPN*-associated muscular dystrophy ^1–3^.

### *snupn* deficiency disrupts RNA processing and muscle niche integrity

To investigate the molecular mechanisms underlying muscle degeneration in MZ *snupn*^⁻/⁻^ larvae, we performed RNA sequencing (RNA-seq) at 6 dpf, a developmental stage at which mutant larvae exhibit clear structural and functional muscle impairment. Principal component analysis (PCA) revealed clear segregation between control and mutant transcriptomes (Fig. S4a), with 602 significantly differentially expressed genes (DEGs) identified, including 288 upregulated and 314 downregulated transcripts (Fig. 4a).

**Figure 4.**
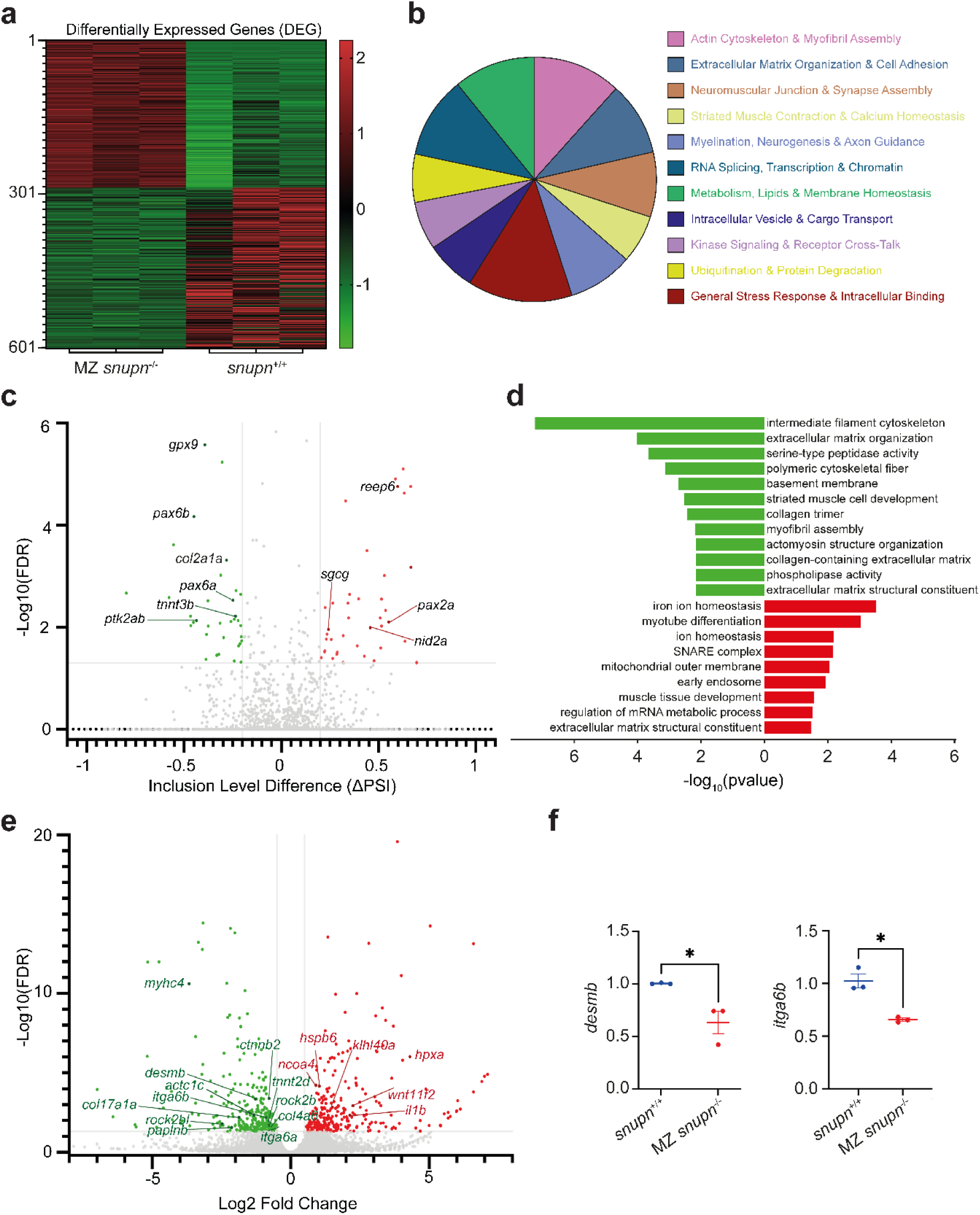
Transcriptomic landscape of MZ *snupn*^−/−^ larvae reveals ECM structural and niche collapse. **a,** Heatmap illustrating 602 differentially expressed genes (DEGs) between control and MZ *snupn*^−/−^ larvae at 6 dpf. n=3 biological replicates per group. **b**, Pie chart depicting the cellular function associations of genes undergone alternative splicing events in MZ snupn^−/−^ larvae. **c,** Volcano plot displaying the inclusion level differences of alternatively spliced genes in MZ snupn^−/−^ larvae. **d,** Gene Ontology (GO) enrichment analysis highlighting the most significantly upregulated (red) and downregulated (green) biological processes in mutants compared to controls. **e,** Volcano plot displaying the distribution of significant DEGs; green and red dots indicate significantly downregulated and upregulated genes, respectively. **f,** qPCR validation of *desmb* and *itga6b* genes shown to be differentially expressed in RNA-seq. Data represent mean ± SEM, n=3 independent experiments; *p=0.02.

Given the role of *SNUPN* in snRNP nuclear import, we first examined alternative splicing and observed extensive defects, predominantly exon skipping and alternative splice-site usage (Fig. S4b). Among the 75 significantly mis-spliced genes, many were associated with neuromuscular organization, extracellular matrix (ECM), and cytoskeletal functions (Fig. 4b), linking defective RNA processing to pathways essential for muscle integrity. Notably, aberrant splicing of the ECM-associated gene *col2a1a* further linked defective RNA processing to ECM organization (Fig. 4c; Fig. S4c).

Consistent with these splicing alterations, transcriptional changes revealed coordinated disruption of muscle structural programs (Fig. 4d; Fig. S4d), with significant downregulation of genes encoding contractile and cytoskeletal components, including *myhc4,* actin genes (*actc1a, actc1c*), *desmb*, and *tnnt2d*, consistent with broad disruption of myofibrillar and cytoskeletal organization (Fig. 4e-f). Notably, *desmb*, the zebrafish orthologue of human *DES*, further connects these structural changes to mechanisms implicated in inherited myopathies ^27^.

Genes associated with ECM organization and cell–matrix adhesion were also altered, including reduced expression of *paplnb*, collagen genes (*col17a1a, col4a6*), and integrin genes (*itga6a, itga6b)* (Fig. 4e-f; Fig. S4d), consistent with impaired ECM organization and muscle fiber–matrix interactions. Alterations in *rock2b* and *rock2bl*, regulators of actomyosin contractility and cell adhesion, further implicate cytoskeletal and cell–matrix organization.

These alterations were accompanied by activation of stress and inflammatory programs, including upregulation of muscle stress markers (*klhl40a, hspb6*), iron homeostasis regulators (*hpxa, ncoa4*), and inflammatory mediators (*il1b*) (Fig. 4d and Fig. S4d). Regenerative signaling was also perturbed, with increased *wnt11f2* and reduced *ctnnb2* (β-catenin), suggesting dysregulation of pathways involved in muscle repair and regeneration (Fig. 4d; Fig. S4d).

Finally, splicing alterations affected key developmental and progenitor-associated regulators, including *pax2a*, *pax6a/b*, the satellite cells–associated receptor *fgfr4*, and focal adhesion kinase *ptk2ab*, pointing to defects in progenitor identity, activation, and niche interaction (Fig. 4c).

Together, these transcriptional and splicing alterations provide a mechanistic framework for the impaired muscle organization observed in *snupn*-deficient larvae, implicating early dysfunction of the myogenic progenitor niche.

### snupn deficiency impairs early myogenic progenitor development

To determine whether late-stage muscle failure is preceded by early defects in the muscle progenitor compartment, we next examined myogenic progenitors at 2 dpf, shortly before the emergence and expansion of Pax7⁺ progenitors that contribute to post-embryonic muscle growth and regeneration ^20^.

Consistent with a potential role in progenitor biology, analysis of the DanioCell dataset revealed that *snupn* is enriched in satellite cells (SCs), supporting a potential role for *snupn* in this compartment (Fig. S5a-e). We first assessed the expression of canonical myogenic progenitor markers (Fig. 5a). Both *pax3a* and *pax7b* were significantly reduced in MZ *snupn*^⁻/⁻^ at 2 dpf (Fig. 5b). In agreement, whole-mount immunofluorescence analysis showed a marked reduction in Pax7⁺ cells, indicating a reduction in the Pax7⁺ muscle progenitor population at this early developmental stage (Fig. 5c).

**Figure 5:**
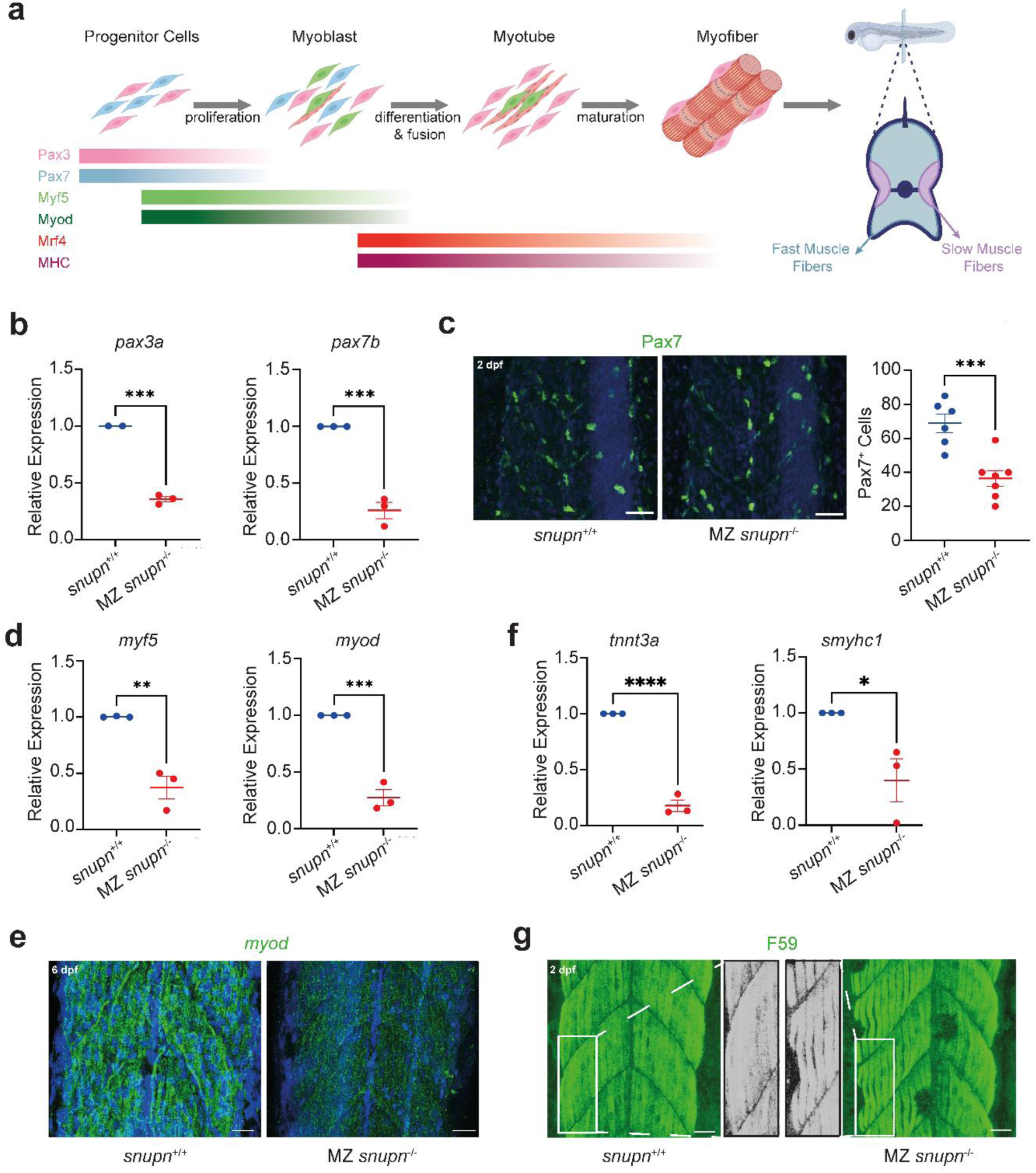
*snupn* deficiency reduces muscle progenitor cells and impairs myogenesis. **a,** Schematic depicting the progression of zebrafish skeletal muscle development highlighting selected genes involved in progenitor specification, myogenic differentiation, and myofiber maturation. **b,** qPCR analysis revealing significant decrease in *pax3a* and *pax7b* mRNA levels in MZ *snupn*^−/−^ mutants compared with WT siblings at 2 dpf. Data represent mean ± SEM, n=3, ***p=0.0038 (left), ***p=0.0005 (right). **c,** Representative immunofluorescence staining and graph showing significant reduced number of Pax7 positive cells in MZ *snupn*^−/−^ mutants compared with WT siblings at 2 dpf. Scale bar: 10 µm. Data represent mean ± SEM, n= 6-7 fish/genotype, ***p=0.0008. **d,** qPCR analysis showing significant reduction of *myf5* and *myod1* transcripts in MZ *snupn*^−/−^ mutants compared with WT siblings at 2 dpf. Data represent mean ± SEM, n=3, **p=0.0036, ***p=0.0005. **e,** Representative HCR image showing reduced *myod1* expression at 6 dpf confirmed. Scale= 25 µm. **f,** qPCR analysis revealing significant reduction in *tnnt3a* and *smyhc1* transcripts levels in MZ *snupn*^−/−^ mutants compared with WT siblings at 2 dpf. Data represent mean ± SEM, n=3, ***p<0.001, *p=0.036). **g,** Representative immunofluorescence staining showing significant reduced level of F59 in MZ *snupn*^−/−^ mutants compared with WT siblings at 2 dpf.

We next examined the activation state of myogenic progenitors by analyzing the early myogenic regulators *myod1* and *myf5*. Both transcripts were significantly decreased in mutants, as shown by quantitative PCR (Fig. 5d). This reduction was further validated by HCR, which confirmed decreased *myod1* expression within muscle tissue (Fig. 5e), supporting impaired myogenic progenitor activation and differentiation.

We then evaluated whether these early progenitor defects impacted downstream differentiation. Expression of muscle fiber differentiation markers, including *tnnt3a* (fast-twitch fibers) and *smyhc1* (slow-twitch fibers), was significantly reduced at 2 dpf in mutant larvae (Fig. 5f). Consistently, F59 immunostaining revealed disrupted and disorganized slow muscle fiber architecture in mutant larvae, indicating defective muscle fiber organization during early muscle development prior to overt functional decline (Fig. 5g). Together, these data demonstrate that *snupn* deficiency impairs the early muscle progenitor compartment, characterized by reduced Pax7⁺ progenitor abundance and defective activation of the myogenic program. These abnormalities precede the later structural and transcriptional disruption of the muscle niche, suggesting that early defects in progenitor expansion and myogenic differentiation may contribute to progressive muscle pathology (Fig. 2 and 6).

**Figure 6:**
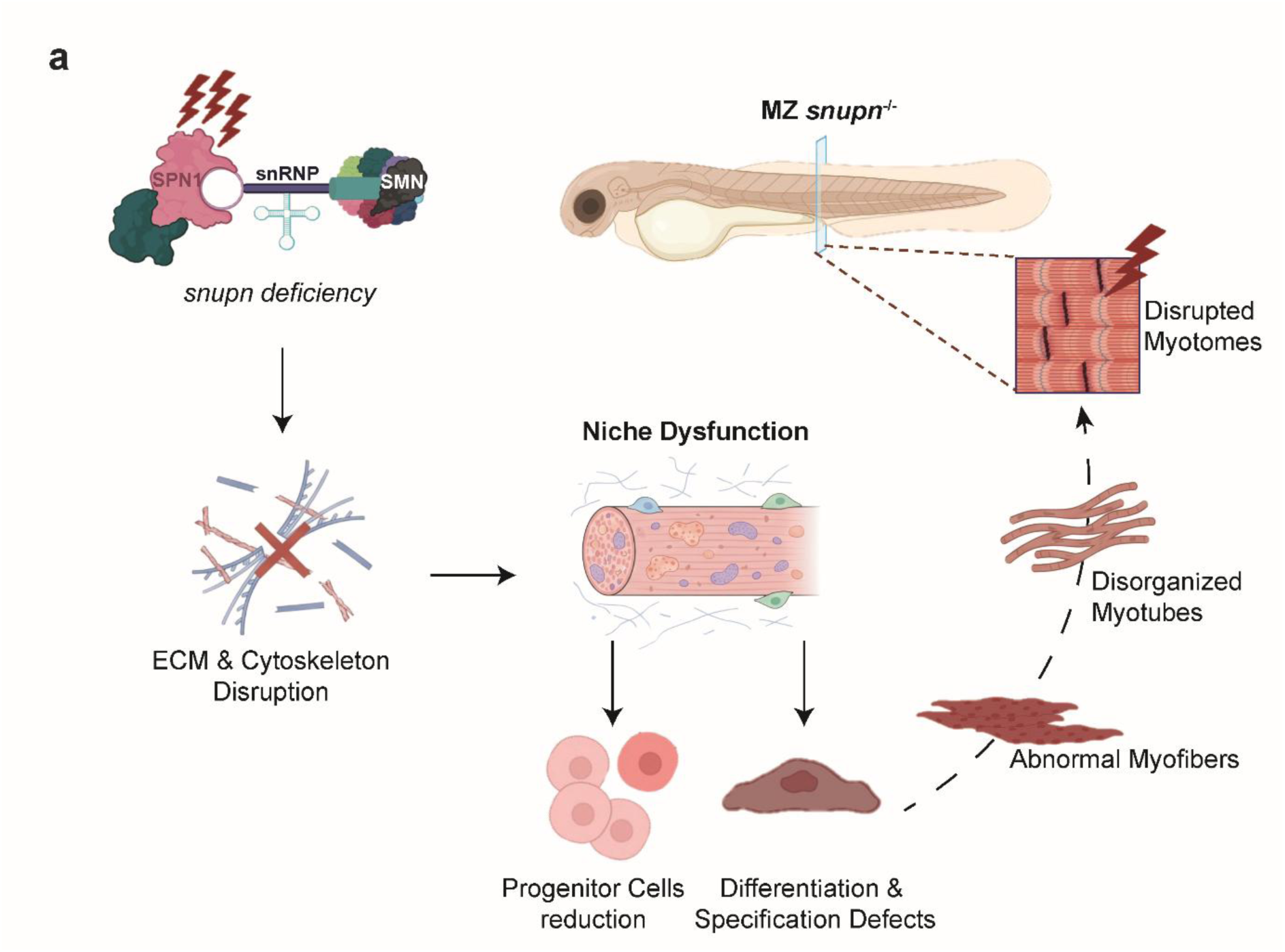
Proposed model of *snupn* dependent muscle pathology. Defective *snupn* leads to disruption of snRNP-mediated RNA processing, resulting in alterations in cytoskeletal organization, ECM integrity, and metabolic pathways. These molecular changes impair muscle niche structure and muscle progenitors function, leading to defective myogenic differentiation and altered muscle fiber type specification. Collectively, these defects contribute to the structural and functional muscle abnormalities observed in the MZ *snupn*^−/−^ mutant line. Created with BioRender.com.

## Discussion

In this study, we demonstrate that the spliceosome-associated nuclear import factor *SNUPN* is required for skeletal muscle homeostasis in a vertebrate model. Using a CRISPR/Cas9-generated zebrafish model, we show that *snupn* deficiency causes progressive structural and functional muscle defects accompanied by widespread transcriptomic dysregulation, early abnormalities in the muscle progenitor compartment, and impaired myogenic differentiation, providing *in vivo* insight into how disruption of *SNUPN*-dependent RNA processing leads to muscle pathology.

Comparison with mammalian models highlights important differences in the consequences of *SNUPN* deficiency. While a knock-in mouse model carrying *SNUPN* variants reproduces cerebellar phenotypes with limited skeletal muscle involvement complete *Snupn* loss is embryonically lethal in mice ^9^. In contrast, MZ *snupn*^⁻/⁻^ zebrafish are viable and develop progressive muscle pathology and severe homozygous *SNUPN* variants in humans are compatible with postnatal survival ^2,3^. These species differences may reflect variation in maternal contribution, developmental compensation, or tissue-specific sensitivity to disrupted RNA processing, raising the possibility that tissue vulnerability to *SNUPN* deficiency is context-dependent. Beyond the muscle phenotype, MZ *snupn*⁻/⁻ zebrafish provides an opportunity to investigate the neurological consequences of *SNUPN* deficiency and may help elucidate the mechanisms contributing to the distinct tissue-specific manifestations observed across species.

Consistent with previous findings in human cells carrying pathogenic *SNUPN* variants, which show extensive splicing defects ^2,3^, *snupn* deficiency in zebrafish caused widespread alternative splicing and transcriptional changes affecting genes involved in sarcomere organization, cytoskeletal integrity, and ECM composition. Importantly, these changes involved ECM and basement membrane components, potentially compromising structural support for muscle fibers and progenitor function and thereby disrupting the muscle niche.

At the cellular level, we observed an early reduction in Pax7⁺ progenitor cells and impaired expression of key myogenic regulators, including *myod* and *myf5*. These defects preceded overt muscle degeneration, indicating that impaired progenitor maintenance and activation are early features of the phenotype. In zebrafish, post-embryonic muscle expansion relies on Pax7⁺ progenitors that proliferate and differentiate into myoblasts, contributing to fiber growth and maintenance ^19,20^. Disruption of progenitor maintenance and activation may therefore contribute to the broad impairment of muscle fiber differentiation observed in *snupn* mutants. Together, the transcriptomic and cellular alterations are consistent with a dystrophic process characterized by disruption of muscle–niche interactions and altered regenerative signaling, rather than a primary developmental defect.

The sensitivity of muscle progenitors to *SNUPN* deficiency may relate to the requirement for precise RNA regulation during transitions between quiescence, activation, proliferation, and differentiation. In murine muscle stem cells, *Snupn* expression is enriched in the quiescent state and decreases upon activation, similar to *Pax7* expression ^28^. Similarly, *snupn* is expressed in zebrafish satellite-like cells in the Zebrahub atlas ^22^, suggesting a conserved association with the muscle progenitor compartment. Other genes involved in muscle integrity and niche function, including *dag1* and *lama2*, show similar expression dynamics ^11,28^. Furthermore, *fgfr4* associated with the quiescent satellite-cell state is reduced upon activation ^28,29^, and its aberrant splicing in *snupn* mutants links altered RNA processing to progenitor and niche regulation. Although correlative, these observations raise the possibility that altered RNA processing compromises the progenitor response to activation and differentiation.

Together, our data support a model in which muscle degeneration arises from a combined defect in both the progenitor compartment and its microenvironment (Fig. 6). Early abnormalities in Pax7⁺ progenitors and myogenic activation are followed by extensive disruption of ECM and basement membrane programs, which may further compromise muscle maintenance and regeneration. This reciprocal interaction between progenitor dysfunction and niche instability could progressively limit the capacity of muscle to compensate for ongoing damage. Defining the specific splicing events that drive these phenotypes, and distinguishing cell-autonomous from niche-mediated effects, will require cell-type-specific models, targeted rescue experiments, and myogenic progenitor cultures.

The *snupn*-deficient zebrafish model provides a tractable system for therapeutic studies, including compound screening and *in vivo* evaluation of candidate interventions. Our findings establish *SNUPN*-dependent RNA processing as a critical determinant of skeletal muscle homeostasis and identify its downstream consequences as potential therapeutic targets. Defining the pathogenic splicing events will be an important step toward developing strategies to restore RNA processing and muscle homeostasis in *SNUPN*-associated disease.

## Acknowledgements

We are grateful to all members of the Department of Medical Genetics, Koç University School of Medicine (KUSoM) for their support and constructive feedback. The authors gratefully acknowledge the Koç University Hospital Department of Pathology for their assistance with histological section preparation and the Koç University Nanofabrication and Nanocharacterization Center for Scientific and Technological Advanced Research (n2STAR) facility for their technical assistance with electron microscopy experiments. This work was conducted as part of the PhD dissertation of the first author, Hilal Pırıl Saraçoğlu, awarded by Koç University in 2026 (#1013986). The authors gratefully acknowledge the services and facilities provided by the Koç University Research Center for Translational Medicine (KUTTAM), particularly the Zebrafish Facility and its staff.

## Author contributions

N.E.B. directed the project, designed and analyzed all the experiments. H.P.S., E.Y., M.N., B.S. and N.N. contributed to zebrafish line generation, including founder screening, genotyping, mutation analysis and sequencing. H.P.S. performed HCR, immunofluorescence staining, qPCR, birefringence assays, and brightfield, histological, and confocal imaging, conducted image analyses and database searches, and prepared the figures. Ş.T. assisted H.P.S. with sample collection for multiple experiments and maintenance of the line. D.N.K. and Em.Y. performed and analyzed behavioral assays. M.N. performed the majority of the omics analyses, with contributions from H.P.S. H.K. and E.Y. initiated the establishment of the zebrafish facility at KUTTAM. N.E.B. and H.P.S. wrote the manuscript with the input of all the co-authors.

## Material and Methods

### CRISPR-mediated zebrafish knockout

Zebrafish were maintained and used in accordance with the Koç University Animal Experiments Local Ethics Committee, Koç University, Turkey (2020-HADYEK-021). A guide RNA targeting exon 3 of *snupn* 5ʹ-CTGGAGCACAGCCGACAGTG-3ʹ was designed. The sgRNA was synthesized using the HighYield T7 sgRNA Synthesis Kit (SpCas9) (Jena Bioscience, Germany) following the manufacturer’s protocol, and the DNA template was digested with DNase Turbo™ DNAse (ThermoFischer, USA). Approximately 75-150 pg of sgRNA and 100-200 pg of Cas9 protein were injected into the yolk of one-cell wild-type (WT) zebrafish embryos. Genotyping was performed by PCR followed by Sanger sequencing. Primer sequences are provided in Supplementary Data 1.

### RNA Isolation and quantitative RT-PCR

Total RNA was extracted from 30-50 zebrafish embryos at the designated developmental stages. RNA from embryos prior to 24 hpf was extracted using TRIzol^TM^ reagent (Invitrogen^TM^, USA), following the manufacturer’s instructions. RNA from 48 and 72 hpf embryos was isolated with the RNeasy Mini Kit (Qiagen) including on-column DNase digestion. cDNA was synthesized from 500-1000 ng of total RNA using the iScript cDNA Synthesis Kit (BioRad, USA). Quantitative RT-PCR (qPCR) was performed with LightCycler^®^ 480 SYBR Green (Roche, Switzerland) on a PikoReal Real-Time PCR System (ThermoScientific, USA). Gene expression levels were normalized to *18S*. Data represent mean ± SEM of biological replicates. Primer sequences are listed in Supplementary Data 1.

### Hybridization Chain Reaction (HCR)

The HCR procedure consisted of four steps: sample preparation, probe hybridization, amplification, and counterstain/imaging (Molecular Instruments, 2025). Buffers and solutions provided by the manufacturer were used according to the supplied protocols (Molecular Instruments, USA). Custom probes targeting *snupn* and *myod* were designed and synthesized by Molecular Instruments.

### Birefringence Assay

Muscle integrity was evaluated using birefringence under polarized light on a Leica DMi8 SP8 Inverted Confocal Microscope (Leica, Germany). 5 dpf larvae were embedded in 1% low melting agarose for imaging, and polarized light settings were applied as described previously.

### Zebrafish Whole Mount Immunofluorescence

Embryos at 48 hpf and larvae at 6 dpf larvae are fixed in 4% paraformaldehyde (PFA) in PBS overnight at +4°C. Samples were dehydrated through a PBS/methanol gradient and stored in 100% methanol at −20°C. Before staining, samples were rehydrated through a reverse gradient of PBS/methanol. Samples were rinsed once with 1xPDT (1xPBST, 0.3% Triton-X, and 1%DMSO) and washed twice on a rocker for 30 minutes at room temperature. Blocking was performed in blocking buffer (1xPBS, 10% heat-inactivated fetal bovine serum, and 2% bovine serum albumin) for 1 hour at room temperature. Primary antibody incubation was carried out overnight at +4°C on a rocker. The next day, samples were washed twice in PDT for 30 minutes each at room temperature, followed by incubation with secondary antibodies for 2 hours at room temperature. After two additional washes in PDT, samples were mounted in 1% low melting agarose on glass slides and imaged using a Leica DMi8 SP8 Inverted Confocal Microscope (Leica, Germany).

### Histological Examination

Zebrafish were euthanized and fixed in 4% PFA overnight at +4°C. Fixed samples were processed by Koç University Hospital Department of Pathology. After 10% formaldehyde fixation, samples were embedded in paraffin using a Tissue-Tek VIP^®^ 6 AI Tissue Processor (Sakura Finetek, USA). Sections (2 µm thick) were stained with hematoxylin and eosin (H&E) using a Tissue-Tek Prisma^®^ Plus Automated Slide Stainer (Sakura Finetek, USA).

### Electron Microscopy

Fish were euthanized on ice. Heads and tails were removed, and each trunk was divided into three segments. Samples were fixed in glutaraldehyde buffer overnight at +4°C, followed by 1 hour of osmium tetroxide treatment. Samples were washed twice in PBS and then treated with uranyl acetate for an hour. After two PBS washes, dehydration was performed through ethanol gradient, followed by two propylene oxide treatments. Samples were embedded in epoxy resin, sectioned at 70 nm thickness, and contrasted with uranyl acetate before imaging with a Hitachi HT7800 transmission electron microscopy (TEM) (Hitachi, Japan).

### Behavioral Analysis

The behavioral test through Light/Dark Transition experiment was conducted in a Zantiks MWP set up using 24-well plates that allow tracking of the animal movement from above. In short, fish were introduced to the well plates and placed in the set up. After a habituation period of 30 minutes, the behavioral protocol started with the dark condition (5 minutes) and then back to the light condition (5 minutes). This was repeated 10 times. After 10 cycles of light on and light off periods, in the last 20 minutes fish were left for resting. The total length of the experiment was 2 hours 30 minutes. The fish were then tracked for their x and y position and the data later analyzed with custom MATLAB scripts.

### RNA Sequencing and Differential Expression Analysis

Total RNA was extracted from triplicate pools of 30–50 6-dpf wild-type (WT) and MZ *snupn*^−/−^ mutant larvae. Library construction (DNBSEQ PE150) and sequencing were performed by BGI (China). Raw sequencing reads were filtered using SOAPnuke (v1.5.6). Clean reads were aligned to the zebrafish reference genome (*Danio rerio*, Ensembl release 105) using HISAT2 (v2.2.1), and transcript quantification was performed using Bowtie2 (v2.5.0) and RSEM (v1.3.1). Differential gene expression analysis between WT and MZ *snupn*^−/−^ mutants was calculated using DESeq2 (v1.40.2), with statistical significance defined as an adjusted P-value (FDR) ≤ 0.05 and an absolute log2 fold change ≥ 0.5. Gene expression levels were quantified as Transcripts Per Million (TPM) and log2-transformed [log2(TPM+1)]. Principal component analysis (PCA) and volcano plots were generated using GraphPad Prism (v10). Heatmaps of differentially expressed genes (DEGs) were constructed from Z-score transformed expression values using hierarchical clustering (Euclidean distance).

### Gene Ontology (GO) Enrichment Analysis

Functional enrichment analysis for DEGs and alternatively spliced genes was performed using the SRPlot platform. For alternative splicing enrichment, an initial list of 97 splice-dysregulated transcripts was filtered to remove redundant distinct exons from the same locus, collapsing the dataset to 84 unique genes. Ten uncharacterized loci lacking established biological process annotations were excluded, yielding a core dataset of 74 mappable genes. GO biological process overrepresentation was calculated using Fisher’s Exact Test against the *Danio rerio* reference background, with P-values adjusted via the Benjamini-Hochberg False Discovery Rate (FDR). Separate enrichment analyses were conducted for all DEGs, as well as upregulated and downregulated subsets, and visualized via bubble plots.

### Alternative Splicing & Visualization

Differential alternative splicing events including skipped exons (SE), retained introns (RI), alternative 3’ and 5’ splice sites (A3’SS/A5’SS), and mutually exclusive exons (MEX) were identified using rMATS (v4.1.2). To qualitatively assess differential exon usage, Sashimi plots were generated directly from aligned, sorted BAM files using the ggsashimi command-line tool. The zebrafish reference annotation (Danio_rerio.GRCz11.105.gtf) was used to map genomic coordinates and overlay transcript models. Intronic regions were graphically compressed, and alpha blending was set to 0.25. Low-confidence splice junctions were filtered using a minimum junction read depth of 10 (-M 10). Biological replicates for WT and MZ *snupn*^−/−^ mutants were plotted independently without merging using a custom color palette (WT in green, MZ *snupn*^−/−^ in red).

### Quantification and statistical analysis

Prism 7 was used to carry out all the statistical tests, unless otherwise stated. Statistical test used for each assay, and number of samples were specified in the Figure legends. The values are presented as mean ± SEM. P-value < 0.05 was considered statistically significant. At least three technical replicates were performed for all assays. All the samples were included, and analyses were performed without blinding.

