## Supplementary data for "Snurportin-1 maintains muscle niche integrity and myogenic progenitor homeostasis"

### *Snurportin-1 links RNA processing to muscle niche integrity and myogenic progenitor homeostasis*

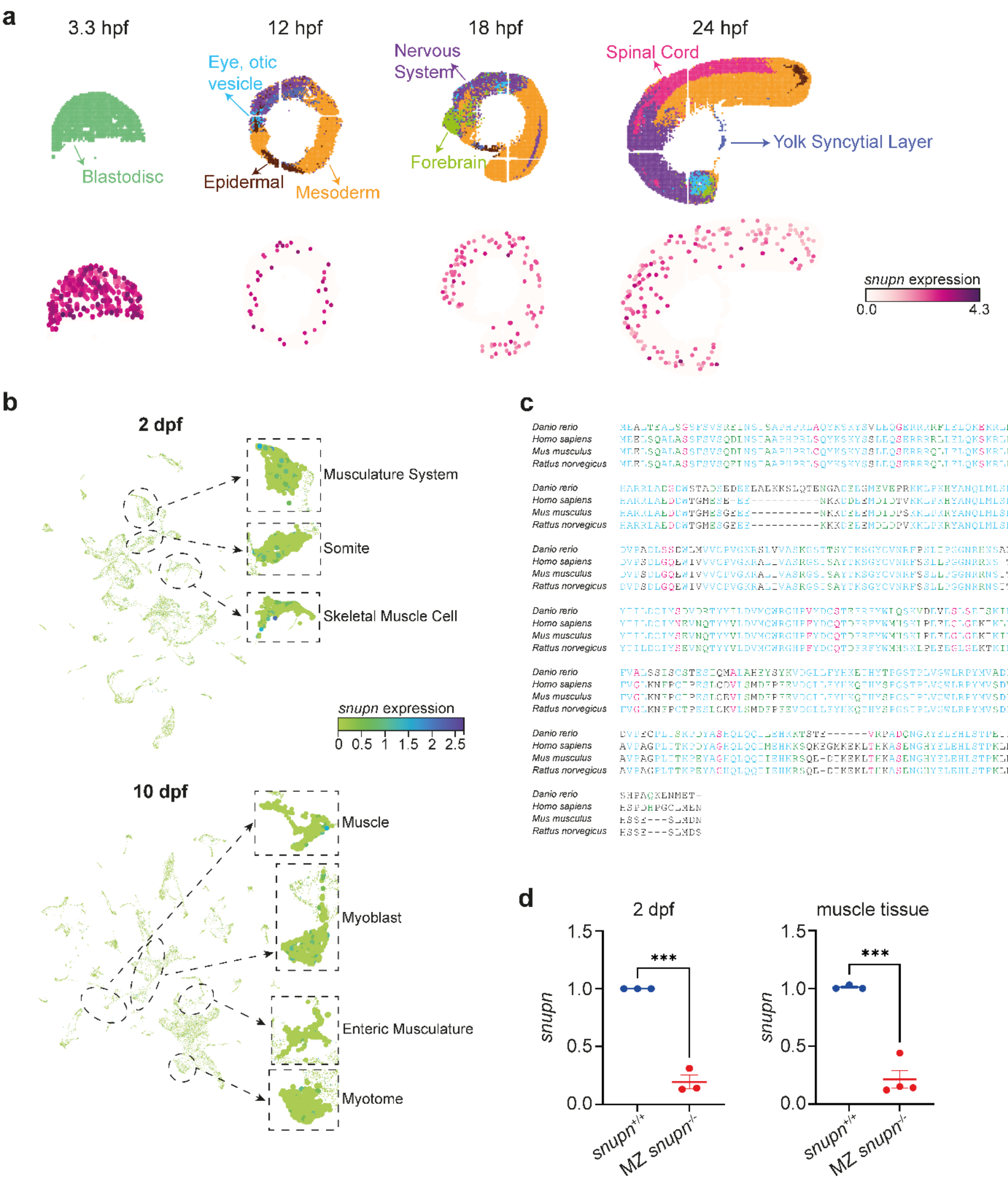


**Supplementary Figure 1. Expression profile and evolutionary conservation of zebrafish *snupn*. a,** Zebrafish Embryogenesis Spatiotemporal Transcriptomic Atlas (ZESTA), showing *snupn* strong expression at the blastomere stage, followed by moderate, widespread expression across all germ layers at 12-, 18- and 24-hours postfertilization (hpf). The main tissues layers are illustrated above and color-coded. **b,** Single-Embryo Single-Cell RNA Sequencing Atlas (Zebrahub) data highlighting *snupn* expression in muscle cell populations at 2- and 10-days post fertilization (dpf). **c,** Phylogenetic analysis showing strong evolutionary conservation of *snupn* across vertebrate species. **d,** qPCR analysis showing reduced *snupn* transcripts in MZ *snupn*$^{-/-}$ embryos at 2 dpf and in adult muscle tissues compared to WT controls. Data represent mean ± SEM, n=4 independent experiments; ***p= 0.0002.

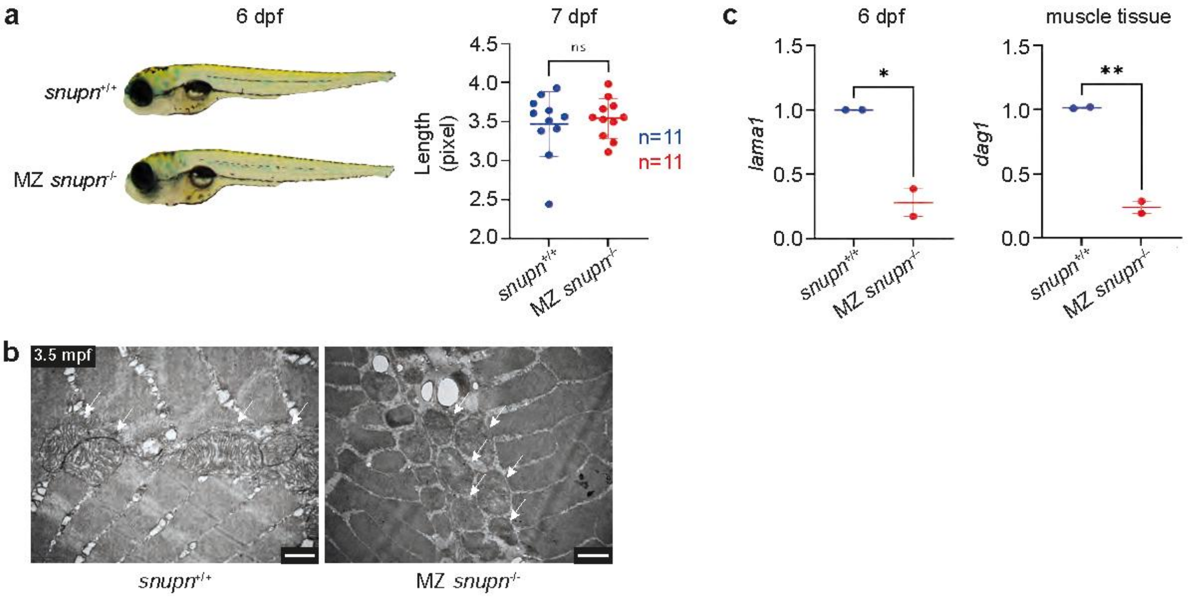


**Supplementary Figure 2: Global morphological analysis of MZ *snupn*$^{-/-}$ mutant larvae. a,** Representative images of 6 dpf WT and MZ *snupn*$^{-/-}$ larvae showing no visible gross morphological differences. Quantification of larval body length at 7 dpf showing no significant difference between WT and MZ *snupn*$^{-/-}$ mutant larvae. n= 11 larvae. Data represent mean ± SEM; ns. **b,** Electron microscopy of adult muscle tissue reveals disrupted mitochondrial morphology in MZ *snupn*$^{-/-}$ compared to WT controls (white arrow). Scale bars: 1 µm. **c,** qPCR analysis showing reduced *lama1* and *dag1* transcripts in MZ *snupn*$^{-/-}$ larvae at 6 dpf and in adult muscle tissues compared to controls. Data represent mean ± SEM, n=2 independent experiments; *p=0.0214; **p=0.0042.

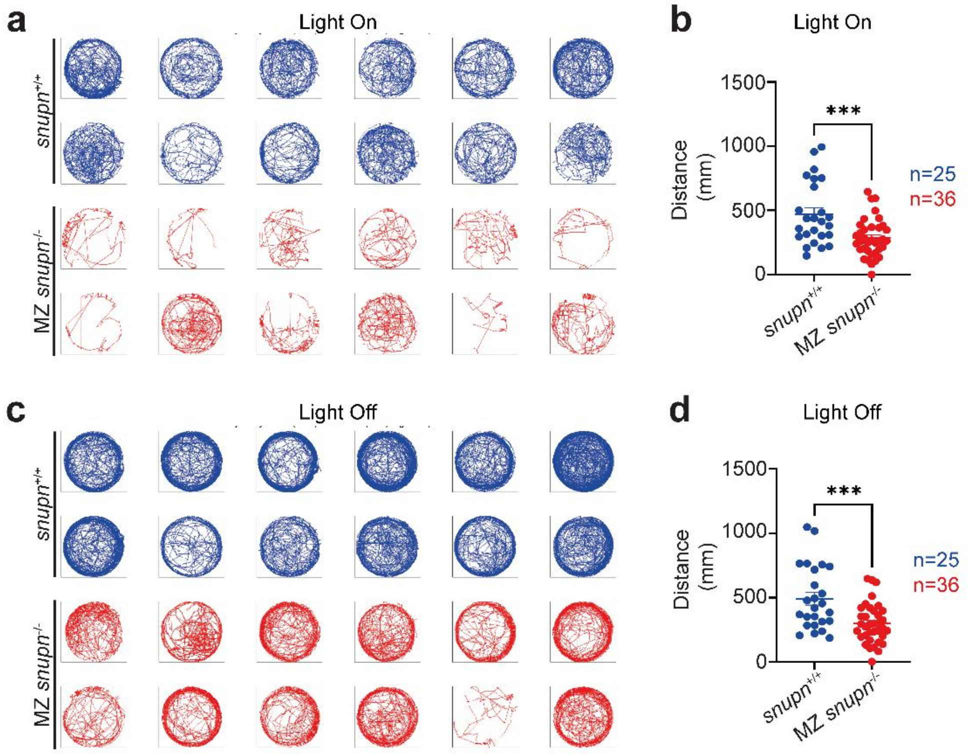


**Supplementary Figure 3: MZ *snupn*$^{-/-}$ deficit of locomotion is independent of light stimuli.** **a**, Representative individual swimming trajectories of 6 dpf larvae, during light on shown in blue for *snupn*$^{+/+}$ and red for MZ *snupn*$^{-/-}$. n=25-36 larvae/genotype. **b**, Graph showing significant reduction in total distance traveled by larvae at 6 dpf covering period of light on. Data represent mean ± SEM, n=25-36 larvae/genotype, ***p=0.0005. **c**, Representative individual swimming trajectories of 6 dpf larvae, during light off shown in blue for *snupn*$^{+/+}$ and red for MZ *snupn*$^{-/-}$. n=25-36 larvae/genotype. **d**, Graph showing significant reduction in total distance traveled by larvae at 6 dpf covering period of light off. Data represent mean ± SEM, n=25-36 larvae/genotype. ***p=0.0004.

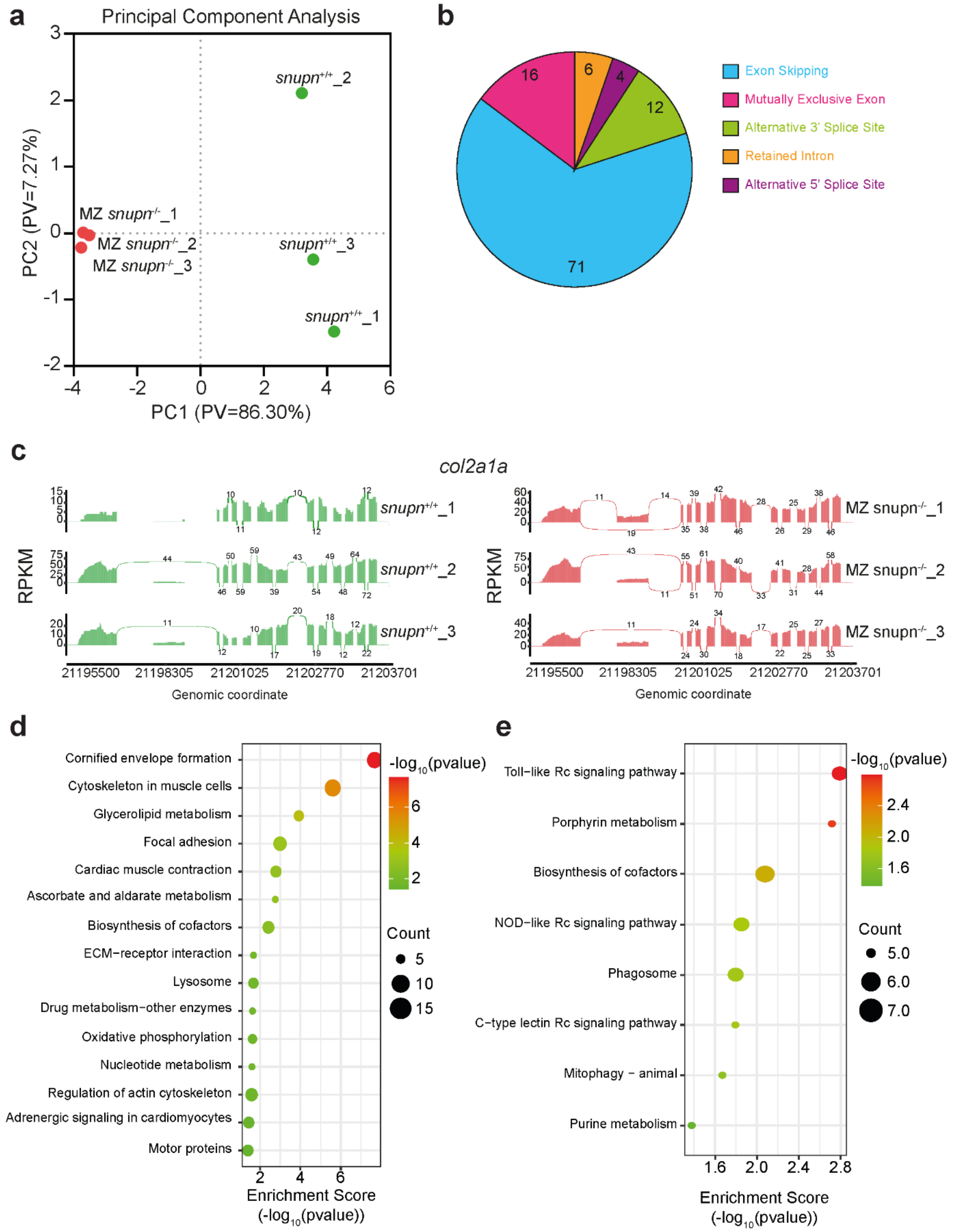


**Supplementary Figure 4: Transcriptomic and alternative splicing alterations in MZ *snupn*$^{-/-}$ larvae.** **a**, Principal component analysis (PCA) showing clear separation between groups of MZ *snupn*$^{-/-}$ (n=3) and control (n=3). **b**, Distribution of alternative splicing (AS) event types in the MZ *snupn*$^{-/-}$ mutant line compared to controls. Relative proportions of major splicing events, including exon skipping (ES), mutually exclusive exon, retained introns (RI), and alternative splice site usage. **c**, Sashimi plots showing significant *col2a1a* gene alterations in the MZ *snupn*$^{-/-}$ mutant line compared to control. Bubble plot of KEGG pathway enrichment analysis of **d,** downregulated and **e**, upregulated genes in MZ *snupn*$^{-/-}$ larvae. Circle size corresponds to the number of DEGs associated with each pathway, and circle color indicates the statistical significance (-log10 p-value, Fisher's exact test), with red representing the highest significance.

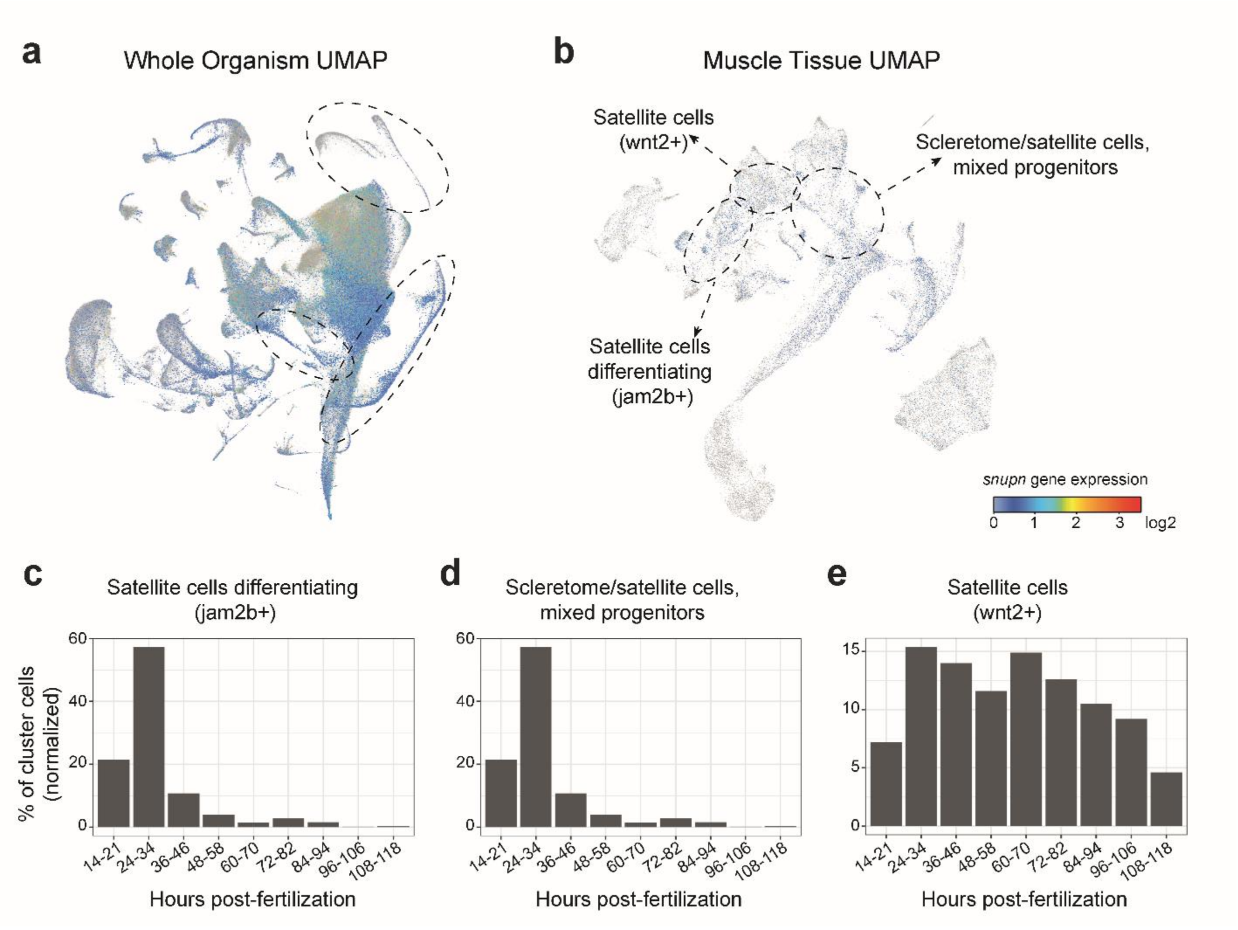


**Supplementary Figure 5: *snupn* expression in the DanioCell single-cell transcriptomic atlas. a,** UMAP showing *snupn* expression across the whole-organism dataset. **b,** Muscle-specific UMAP showing enrichment of *snupn*-expressing cells in muscle cell populations. Color intensity indicates relative gene expression, ranging from low (blue) to high (yellow-red). Developmental dynamics of *snupn* expression in **c,** satellite cell differentiating (jam2b+), **d,** scleretome/satellite cell mixed progenitors, and **e,** satellite cells (wnt2+) populations. Bar plots indicate the percentage of cells expressing *snupn* within each cluster at different developmental time points. Data retrieved from the DanioCell single-cell RNA-sequencing database.

**Supplementary Data 1**

| Gene | Forward Primer | Revere Primer | Method |
|---|---|---|---|
| *snupn* | TGGAGCTGCAGAAAGAGTGA | CCGTTTTGTTCTGTTTACGCA | Sequencing |
| *snupn* | AGAGGACACTCTGGCACATC | ACAAGGAGACGCACTACACC | qPCR |
| *desmb* | ATGCAAGAGACCCAAGTCCA | TCTTGCTCACAGCCTGGTTA | qPCR |
| *itga6b* | ACATTCACAACCCCACCAGAA | TACTCGCTGGCATGACTGAC | qPCR |
| *pax3a* | CTCTGCCATGTCGAGTCTACAG | GAGGCCGTTGCTGATGGAG | qPCR |
| *pax7b* | GGTTCAGCAATCGTCGAGC | CTGGACGCTGATTGGCTCATG | qPCR |
| *myf5* | AACCGGGCCATTGTCTCC | TGCCTCAAAGGCGTGATTG | qPCR |
| *myod* | ACGCCATTAGTTATATCGAGTCT | CTGTCATAGCTGTTCCGTCT | qPCR |
| *tnnt3a* | ACGTCACAACAAGGATACACTTGAGC | CTCTGCTGCTCTGCCCTCTC | qPCR |
| *smyhc1* | TGAAGAGGCTGAGGAACAGG | AGAACCAGTACTTGAACATGGC | qPCR |
| *18S* | TCGCTAGTTGGCATCGTTTATG | CGGAGGTTCGAAGACGATCA | qPCR |
